# miR-10b Deficiency Affords Atherosclerosis Resistance

**DOI:** 10.1101/248641

**Authors:** Masako Nakahara, Norihiko Kobayashi, Masako Oka, Kenta Nakano, Tadashi Okamura, Akira Yuo, Kumiko Saeki

**Affiliations:** Department of Disease Control, Research Institute, National Center for Global Health and Medicine, Tokyo 162-8655, Japan; Department of Laboratory Animal Medicine, Research Institute, National Center for Global Health and Medicine, Tokyo, Japan; Department of Infectious Diseases, Section of Animal Models, Department of Infectious Diseases, Research Institute, National Center for Global Health and Medicine, Tokyo162-8655, Japan.

## Abstract

Human vascular endothelial cells (ECs) are categorized into two groups; pro-stenotic (Type-I) and anti-stenotic (Type-II) ECs, and one of the master genes for a stress-induced “Type-II-to-Type-I” degeneration is Regulator of G-protein signaling 5 (RGS5). Here we show that miR-10b is a crucial downstream mediator in RGS5-dependent degeneration. We also demonstrated the miR-10b^high^ Type-I EC exosome has a trans effect which suppresses anti-proliferative abilities of Type-II ECs. Moreover, we found miR-10b-deficient mice showed a resistance to experimental atherosclerosis, where high-fat-high-cholesterol-diet-fed mice were subjected to partial carotid ligation. Furthermore, we determined the key target of miR-10b was Latent transforming growth factor-β binding protein 1 (LTBP1), which is a regulator of TGF-β signaling. Compatible with a commonly accepted view that TGF-β creates the major growth-inhibitory signal against vascular smooth muscle cells, TGF-β inhibitor treatments abolished anti-proliferative functions of Type-II ECs. Therefore, RGS5/miR-10b/LTBP1/TGF-β axis plays a leading role in quality control of ECs.

## Introduction

Ischemic heart disease and brain stroke are the top two causes of death in the world. The major pathological basis for these diseases is atherosclerosis, where hardening and narrowing (i.e. stenosis) of the arteries causes insufficient blood supply, and thus, impairs the functions of target organs/tissues including the heart, brain, lung and kidneys. Atherosclerosis begins with damage to vascular endothelial cells (ECs), which then leads to the formation of neointima by triggering the hyper-proliferation of vascular smooth muscle cells (SMCs). In addition to atherosclerosis, other pathological situations also induce stenosis of the arteries. For example, patients with systemic sclerosis often suffer from fatal ischemic organ failures due to serious stenosis of the arteries in the lungs and kidneys, where onion skin–like intimal thickening is observed *via* hyper-proliferation of SMCs. Simple mechanical damage to ECs even induces severe stenosis of the arteries; for example, restenosis after stent placement therapy in humans and wire injury operation in mice as its experimental model.

The common pathological feature of arterial stenosis is the hyper-proliferation of SMCs. Although the majority of SMCs reside within the tunica media of the arteries, a small number of them locate within the tunica intima underneath vascular endothelial cells (ECs). When ECs are damaged or removed, SMCs in tunica media start to migrate to tunica intima, where they proliferate without control. The hyper-proliferated SMCs create neointima, which narrows the arterial lumen and reduces blood flow. Although there had been a longstanding dispute over the involvement of ECs in the etiology of ischemia, the controversy has been settled by a series of integrated studies using a wide variety of commercially available primary cultured human ECs and multiple lines of human pluripotent stem cell-derived ECs(1–3). In those studies, it was shown that human ECs are classified into two types according to their effects on the proliferation of SMCs. While commercially available primary cultured ECs are exclusively as pro-proliferative ECs (Type-I)(1), freshly prepared ECs from human embryonic stem (ES) cells are classified as anti-proliferative ECs (Type-II). In addition, these Type-II ECs are converted into Type-I ECs after repetitive subcultures or oxidative stresses(1). The master gene for phenotype conversion from anti-proliferative Type-II into pro-proliferative Type-I ECs is Regulator of G-protein signaling 5 (RGS5)(1): freshly prepared ECs are negative for RGS5 expression while various kinds of stresses including oxidative stress and repetitive subcultures induce RGS5 expressions, and therefore, RGS5 shuts down the anti-proliferative functions of Type-II ECs. By contrast, RGS5 protein expressions are significantly up-regulated in ECs of pathological vessels as observed in the cases of hypertension(1), arteriosclerosis(1), systemic lupus erythematosus(1) and scleroderma(4, 5). Furthermore, transplantation experiments verified the “pro-stenotic” and “anti-stenotic” properties of Type-I and Type-II ECs, respectively(6). In Figure 1A, the concept for Type-I *versus* type-II categorization of ECs is summarized. However, the crucial molecules that are involved in the regulation of Type-I/Type-II phenotypes of ECs at the downstream of RGS5 remain elusive.

**Figure 1.**
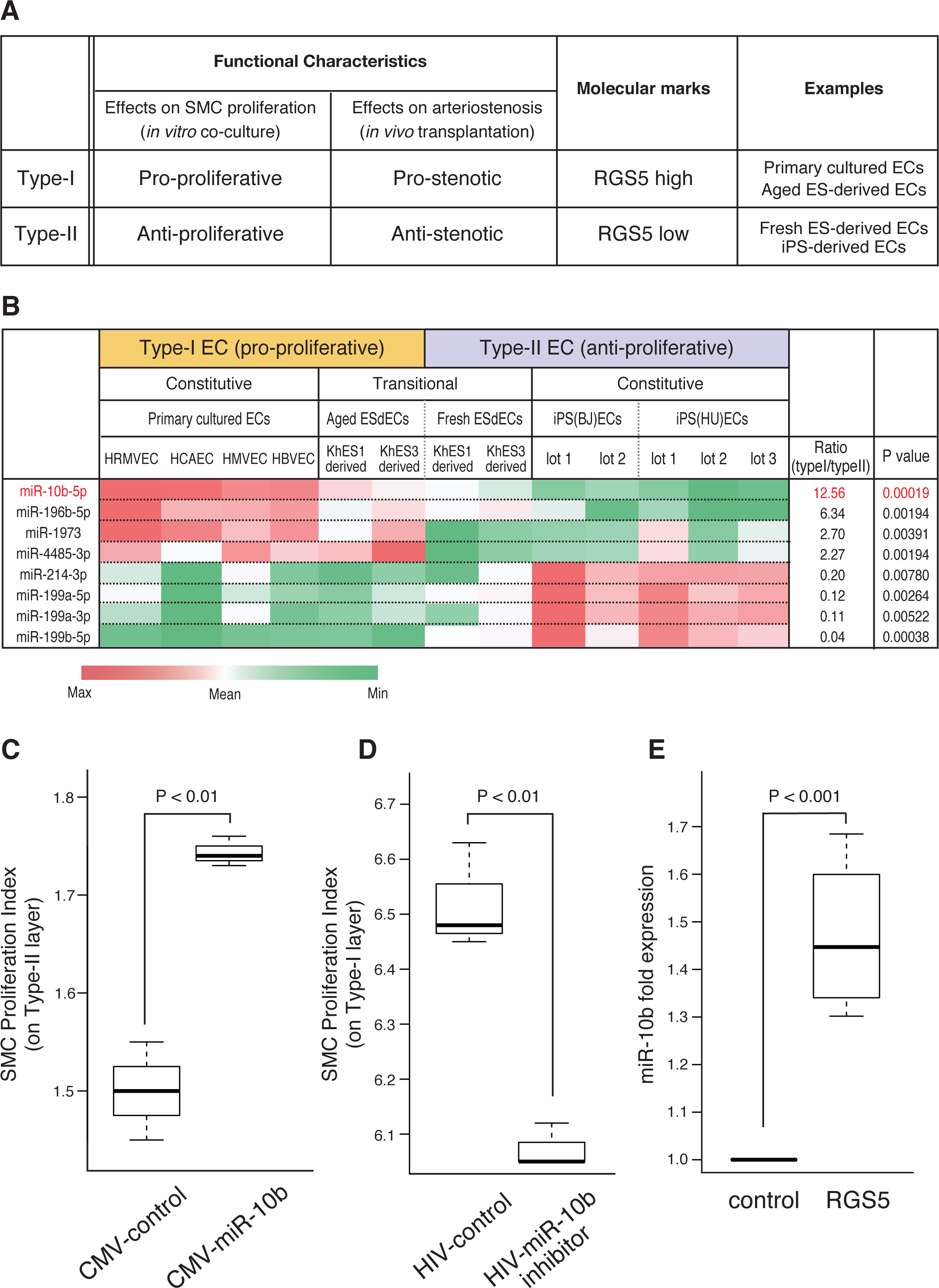
Identification miR-10b as a downstream effector of RGS5. **A.** A brief summary for the concept for Type-I *versus* Type-II categorization of ECs. **B.** Results of the microarray analyses were shown, where the expression profile of each miRNA among various lines of human ECs was demonstrated by an independent heat map. Data regarding miRNAs that showed significantly different (P < 0.01) expression profiles between Type-I EC and Type-II EC groups were shown. Note that the expression profile of miR-10b, which has the smallest P value, shows a precise correlation with Type-I/Type-II phenotyping of ECs. **C.** Proliferation indexes of SMCs that were cultured on the layer of Type-II ECs (ESdEC at early passages), which were transfected with either an empty plasmid (CMV-control) or a miR-10b-expressing vector plasmid (CMV-miR-10b). N=3. **D**. Proliferation indexes of SMCs that were cultured on the layer of Type-I ECs (ESdEC at late passages), which were transfected with either an empty HIV vector (HIV-control) or an miR-10b inhibitor-expressing HIV vector (HIV-miR-10b inhibitor). N=3. **E.** Fold increments in miR-10b expressions in either an empty plasmid (control) or an RGS5-expressing plasmid (RGS5) into Type-II ECs (ESdEC at early passages). N=4. Abbreviations: ESdEC, human ES cell-derived ECs; iPS(BJ)EC, ECs generated from human BJ fibroblast-derived iPS cells; iPS(HU)EC, ECs generated from HUVEC-derived iPS cells.

It is widely accepted that transforming growth factor β (TGF-β) transmits a robust and potent inhibitory signal against SMC proliferation(7–11). Although the expression level of TGF-β defines the strength of growth inhibitory potential in most cases, the other factors also determine the efficacy of TGF-β signaling. For example, Latent transforming growth factor-β binding protein 1 (LTBP1) plays an essential role in the induction of TGF-β mediated quiescence of hematopoietic stem cells (HSCs)(12). Therefore, expression levels as well as activities of TGF-β and LTBP1 should be precisely evaluated to understand the molecular mechanism for Type-I/Type-II quality control of ECs.

Here we showed that miR-10b, which is known to be involved in malignant progression of solid tumors(13, 14), is a crucial downstream mediator of RGS5-dependent “Type-II-to-Type-I” phenotype conversion of ECs. We also showed miR-10b reduces the expression level of LTBP1 protein, thus could suppress the anti-proliferative ability of Type-II ECs. We further demonstrated *in vivo* relevance of miR-10b in the pathophysiology of atherosclerosis by showing that miR-10b knockout mice become resistance to the development of atherosclerosis. Possible link between vascular injuries and malignant progression of tumors is also discussed.

## RESULTS

### miR-10b is the crucial downstream mediator for RGS5-dependent EC phenotyping

To understand the molecular events involved in phenotype regulation of ECs at the downstream of RGS5, we first examined the expression profile of TGF-β. Although TGF-β is known as the major cytokine to suppress the proliferation of suppression of SMCs(7–11), TGF-β mRNA as well as TGF-β protein did not show Type-I/Type-II categorization-dependent expression profiles (Supplemental Figure S1A, S1B). Moreover, we found no correlations between the LTBP1 mRNA expression profile and Type-I/Type-II categorization (Supplemental Figure S2). Therefore, the proliferation of SMCs is regulated by other mechanisms than mRNA expression-regulating system for TGF-β and LTBP1.

Since our previous microarray analyses using exon probes (GEO Accession ID: GSE60999 and GSE61000) revealed that RGS5 was the only protein-coding gene that showed EC categorization-dependent expression profiles(1), we hypothesized that certain non-coding RNAs would be involved in regulating Type-I/Type-II quality control of ECs. From this standpoint, we performed miRNA array analyses using various lines of Type-I and Type-II human ECs. We obtained eight candidate miRNAs, which showed statistically significant differences (P < 0.01) in their expression profiles between the two groups (Figure 1B). They consisted of hsa-miR-10b, hsa-miR-196b-5p, hsa-miR-1973, hsa-miR-4485-3p, which showed Type-I-dominant expression patterns, and hsa-miR-199b-5p, hsa-miR-199a-5p, hsa-miR-199a-3p, hsa-miR-214-3p, which showed Type-II-dominant expression patters. Among these candidates, we focused on has-miR-10b-5p (hereinafter referred to as miR-10b) since its expression profile exactly coincided with the mode of Type-I/Type-II categorization as supported by the lowest P value (P = 0.000186). To verify whether miR-10b was involved in the regulation of Type-I/Type-II quality control of ECs at the downstream of RGS5, we examined the effects of overexpression and knockdown of miR-10b. We showed the enforced expression of miR-10b in Type-II ECs suppressed their anti-proliferative function (Figure 1C), while treatments with a miR-10b inhibitor(15) deteriorated pro-proliferative capacities of Type-I ECs (Figure 1D). We also confirmed that enforced expressions of RGS5 in Type-I EC induced up-regulation of miR-10b (Figure 1E).

Therefore, miR-10b is a crucial downstream mediator in RGS5-dependent Type-I/Type-II quality control of ECs.

### Stress-dependent induction of miR-10b in ECs

Since oxidative stresses up-regulate endothelial RGS5 expressions(1), we examined the effects of various reactive oxygen species (ROS)-producing vasotoxic stimuli on the expression of miR-10b. We found that γ-irradiation(16) (Figure 2A), hyperglycemic stresses(17, 18) (Figure 2B), azidothymidine (AZT) treatments(19) (Figure 2C) induced the expressions of miR-10b, indicating that vasotoxic ROS is an important inducer of miR-10b expressions in ECs.

**Figure 2.**
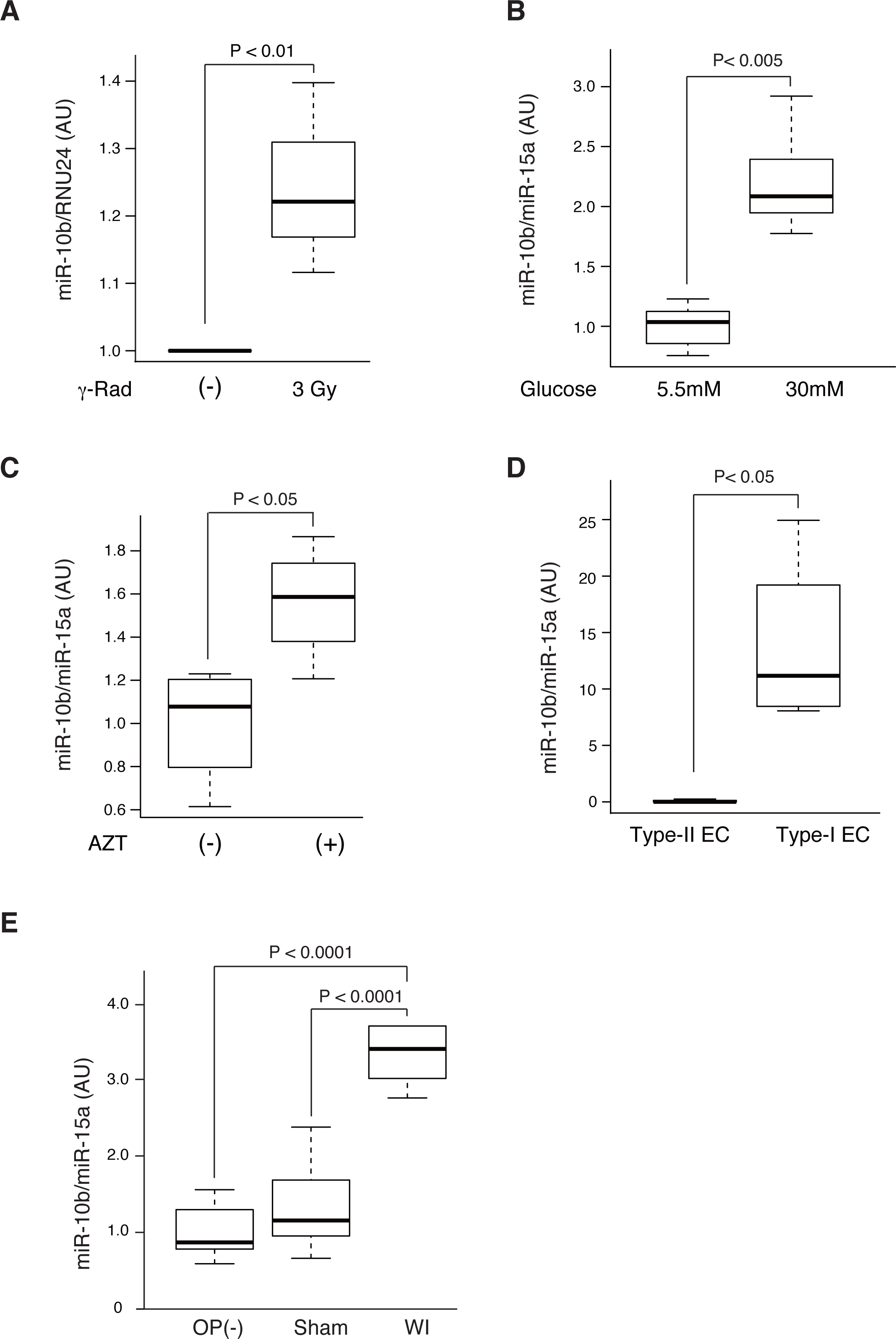
Effects of various noxious stimuli on miR-10b expressions. Expressions of miR-10b were evaluated by qRT-PCR after various toxic stimulations. **A.** miR-10b expressions in HUVEC treated with or without γ-irradiation (5 Gy x 2/week, Day 8) were measured. N=3. **B.** miR-10b expressions in HUVEC cultured under euglycemic (5.5 mM glucose) or hyperglycemic (30 mM glucose) for 4 days were measured N=5. **C.** miR-10b expressions in ES(KhES3)-derived ECs treated with or without 50 μM AZT treatments for one day were measured. N=4. **D**. miR-10b expressions in exosomes collected from the supernatant of human iPS-derived ECs (Type-II) or that of HUVEC (Type-I) were measured. N=4. **E**. miR-10b expressions in blood plasma exosomes of mice without operations (OP(-)), those of mice with sham operations (Sham) and those of mice with wire injury operations (WI) were measures. N=7~11.

It is well known that miRNAs are incorporated into exosomes and transmit various biological effects to the target cells. Therefore, we collected exosomes from Type-I and Type-II ECs (Supplemental Figure S3) and examined the expression levels of miR-10b. We found that Type-I EC’s exosomes contained higher amounts of miR-10b than Type-II EC’s one (Figure 2D), consistently with the finding that Type-I ECs expressed higher levels of intracellular miR-10b (Figure 1B). To verify *in vivo* relevance of the expression of exosomal miR-10b in the context of vascular stresses/damages, we performed wire injury (WI) operations in murine femoral arteries and measured the levels of miR-10b expressions within blood plasma. We found that miR-10b expression levels in circulating blood exosomes were up-regulated after vascular damages (Figure 2E).

Therefore, miR-10b expression levels are up-regulated under EC-damaging circumstances both *in vitro* and *in vivo*.

### miR-10b knockout mice are resistant to atherosclerosis development

To further assess pathological relevance of up-regulated miR-10b expressions with vascular damages, we generated miR-10b knockout (KO) mice (Supplemental Figure S4). To evaluate the involvement of miR-10b in the development of atherosclerosis, wild type (WT) and two lines of miR-10b KO mice were subjected to experimental atherosclerosis. Five-week-old mice, which were fed by high-fat, high-cholesterol diet supplemented with cholic acid, were subjected to partial carotid ligation(20). After three weeks, the conditions of treated and non-treated carotid arteries were examined by ultrasonography. We found that treated arteries of WT mice exclusively showed stenotic features (Figure 3A upper right, white circle) with significantly reduced motilities reflecting sclerotic changes of the vascular walls (Figures 3B upper and Figure 3C). On the other hand, treated arteries of miR-10b KO mice lacked obvious features of stenosis (Figure 3A, lower) and the vascular walls showed higher motilities (Figures 3B lower and Figure 3C). Histological examinations further confirmed that treated arteries of WT mice underwent severe atherosclerosis with nearly complete stenosis (Figure 3D, right panels, White arrow), whereas the lumens of the treated arteries of miR-10b KO mice remained open (Figure 3D, right panel, blue arrow head). Altogether, these findings indicate that miR-10b KO mice become resistant to atherosclerosis development.

**Figure 3.**
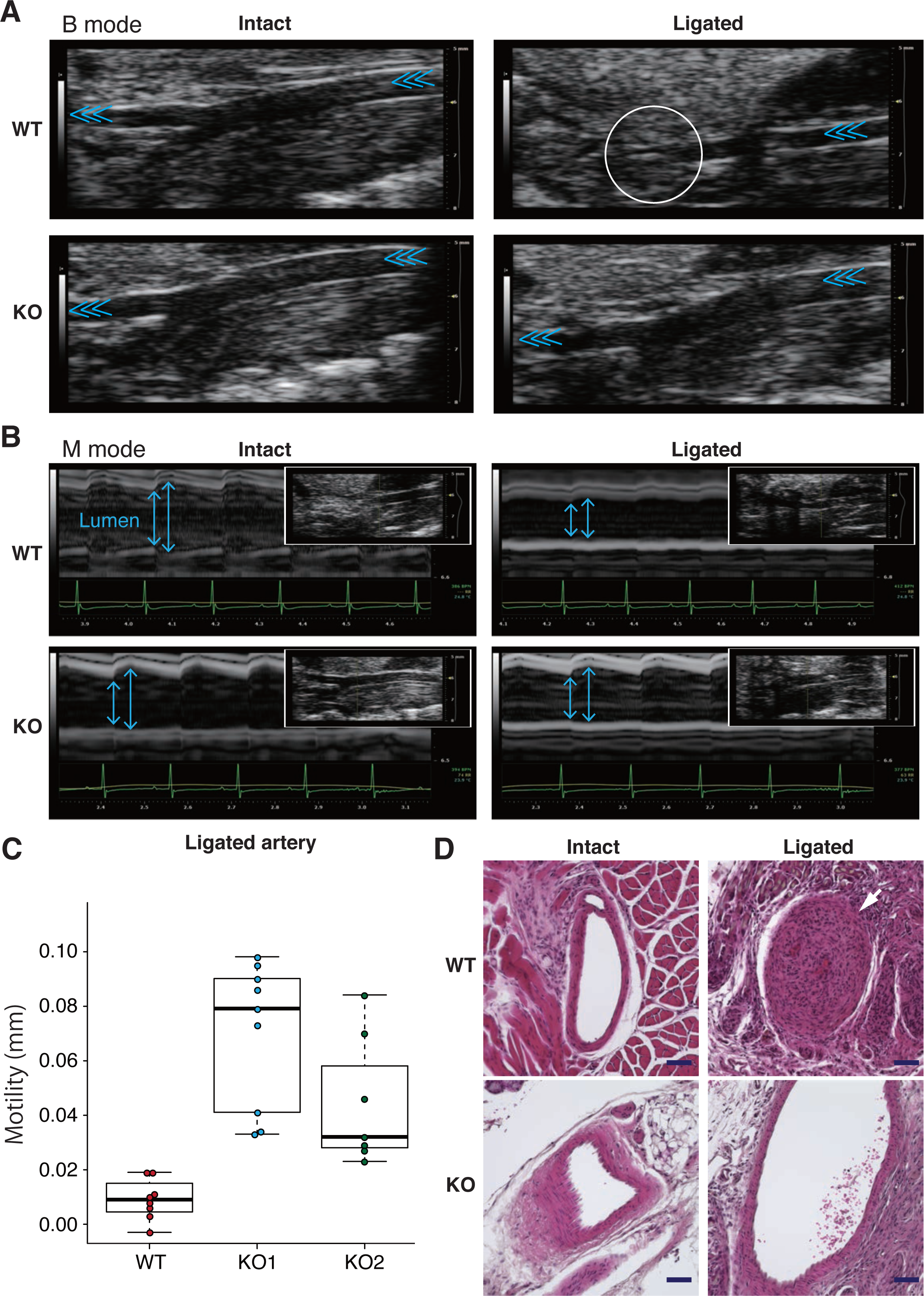
Experimental atherosclerosis. Wild type mice (WT) and miR-10b knock out mice (KO) were subjected to experimental atherosclerosis. Mice were fed by high-fat-high-cholesterol diet and the three branches (external carotid artery, internal carotid artery, occipital artery) out of the four branches of the left common carotid artery were ligated. After three weeks, the conditions of the common carotid arteries were examined. **A and B.** Results of ultrasonically examinations of the carotid arteries were shown by B mode (A) and M mode (B). Light blue triangles in (A) indicate the blood flow in the carotid arteries. A white circle in the right upper panel in (A) indicates the area with stenosis due to the development of atherosclerosis. **C**. Motility of the walls of carotid arteries in (B) were measured and statistically analyzed (WT: n=8, KO1: n=9, KO2: n=7). **D**. Typical findings of the histology of the intact (right) and ligated (left) carotid arteries of wide type (WT) and miR-10b KO (KO) mice were shown. Tissue slices were stained by hematoxylin and eosin staining method. Scale= 50 μm.

### The target of miR-10b is LTBP1 in Type-II EC-mediated growth suppression of SMCs

To identify the target of miR-10b in Type-I/Type-II quality control of ECs, we searched the candidate target mRNAs by using miRDB software(21, 22). As shown in Supplemental Figure S5, we obtained 232 candidates as a target mRNA for miR-10b. Among these candidates, we focused on LTBP1 (Figure 4A) since it is known to play an essential role in transducing TGB-β-dependent growth-inhibitory signaling in HSC niche(12). TGF-β is secreted in a latent form and deposited to extracellular matrix *via* LTBP1, thus making a latent TGF-β/LTBP1 complex. When latent TGF-β is cleaved by specific proteases such as αV Integrins, active TGF-β is generated and released from the complex. Therefore, LTBP1 provides an effective platform for a rapid increment in the local concentration of active TGF-β at the site of action, serving as the key factor in the biological effects of TGF-β in certain cases(12, 23, 24). Hence, we examined the expressions of LTBP1 protein by Western blotting and found that its expression profile precisely coincided with Type-I/Type-II categorization of ECs: Primary cultured ECs with high miR-10b expressions (Type-I) and iPS/ES-derived ECs with low miR-10b expressions (Type-II) were negative and positive for LTBP1 protein expressions, respectively (Figure 4B, left and middle) and miR-10b^low^ ES-derived ECs at early passages (Type-II) and miR-10b^high^ ES-derived ECs at later passages (Type-I) were positive and negative for LTBP1 protein expressions, respectively (Figure 4B, right). In addition, an enforced expression miR-10b in Type-II ECs (Figure 4C) and a transfection of miR-10b inhibitor-expressing vector into Type-I ECs (Figure 4D) resulted in reduction and recovery of LTBP1 protein expressions, respectively.

**Figure 4.**
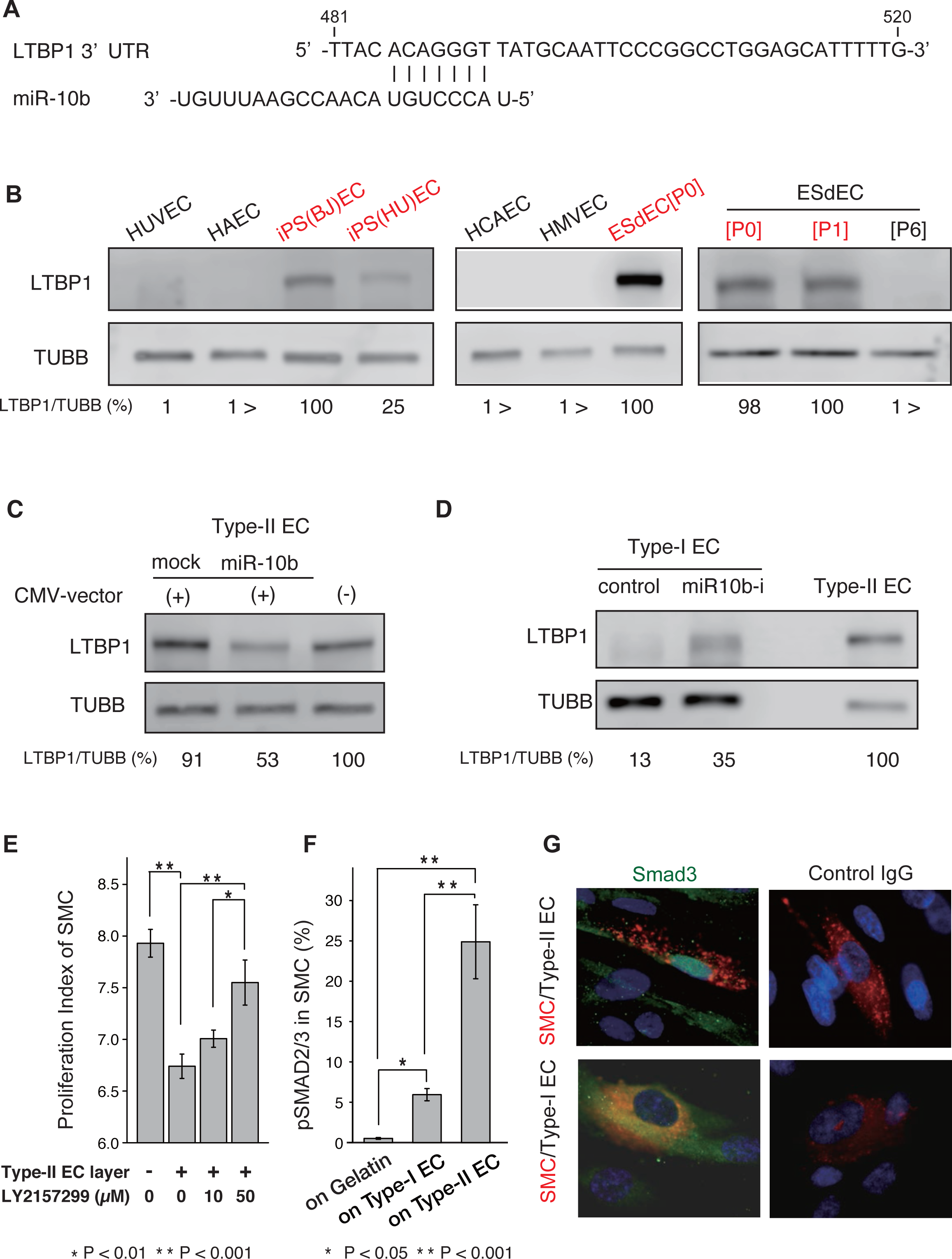
LTBP1 as a target of miR-10. **A.** Nucleotide sequences of miR-10b and those of its target in 3’ UTR in LTBP1. **B.** Western blotting for LTBP1 and β-tubulin (TUBB) using various human ECs that showed Type-I phenotypes (HUVEC, HAEC, HCAEC, HMVEC and ESdEC[P6]) and those with Type-II phenotypes (iPS(BJ)EC, iPS(HU)EC, ESdEC[P0] and ESdEC[P1]) as reported previously^1^. **C**. Type-II ECs that were transfected with an empty vector (CMV-vector (+), mock) or an miR-10b expression vector (CMV-vector (+), miR-10b), and Type-II ECs without transfection (CMV-vector (-)) were subjected to Western blotting for using an anti-LTBP1 body or an anti-β-tubulin (TUBB) antibody. **D**. Western blotting for LTBP1 and β-tubulin (TUBB) proteins using Type-I ECs that were transfected with a control HIV vector (control) or an miR-10b inhibitor-expressing HIV vector (miR-10bi) together with Western blotting using Type-II ECs without transfection were shown as indicated. **E.** Proliferation indexes of SMCs that were co-cultured with Type-II ECs in the absence or the presence of increasing concentrations of LY2157299, a TGF-β signaling inhibitor, as indicated. N=3. **F and G**. SMCs were co-cultured with Type-I or Type-II ECs and the percentages of phosphorylated SMAD2/3-positive cells in were calculated by flow cytometry (F) and nuclear localization of SMAD3 in PKH-26 (red)-stained SMCs was estimated by immunostaining studies with an anti-smad3 antibody (green) along with nuclear counterstaining with DAPI (blue) (G). Full-length blots are presented in Supplementary information. Abbreviations: ESdEC[P6], human ES cell-derived ECs at passage 6; ESdEC[P0], human ES cell-derived ECs at passage 0; ESdEC[P1], human ES cell-derived ECs at passage 1.

We also examined protein expressions of αV Integrins in Type-I and Type-II ECs. The expressions of αV integrin, β5 integrin and β8 integrin were detected in a broad spectrum of human ECs regardless of Type-I/Type-II categorization of ECs (Supplemental Figure S6A), indicating that active TGF-β is readily generated in proportion to the expression levels of LTBP1 protein. In other words, the expression of LTBP1 protein *per se* is the critical determinant for the efficient transmission of TGF-β signaling. In addition, an enforced expression of miR-10b (Figure 4C) or knockdown of miR-10b expression (Figure 4D) impaired or restored LTPB1 protein expressions, respectively. By contrast, the expression profile of LTBP2 protein, which does not bind latent TGF-β(25), was not correlated with Type-I/Type-II categorization of ECs (Supplemental Figure S6B). On the other hand, LTBP3, which could bind latent TGF-β(25), and LTBP4, which scarcely bind latent TGF-β either(25, 26), showed Type-I/Type-II categorization-associated expression profiles (Supplemental Figure S6B). However, an enforced expression of miR-10b as well as knockdown of miR-10b expression did not affect the expression of either LTBP3 or LTBP4 protein (Supplemental Figure S6C). These findings indicate that miR-10b specifically targets the expression of LTBP1 protein.

To verify the involvement of TGF-β-dependent signals in Type-II EC-mediated growth suppression of SMCs, we inhibited the TGF-β signaling in Type-II EC-SMC interactions. When we added LY2157299, a selective small molecule TGF-β receptor I kinase, to the EC-SMC co-culture system, anti-proliferative capacities of Type-II ECs were reduced in a dose dependent manner (Figure 4E). Hence, TGF-β signaling was involved in executing of the anti-proliferative function of Type-II ECs against SMCs. Flow cytometric analyses further demonstrated that the number of SMCs with phosphorylated SMAD2/3, which are the crucial downstream effectors in the intracellular signal transduction after TGF-β receptor activations, was up-regulated by co-culture with Type-II ECs (Figure 4F). Moreover, nuclear localized SMAD3 in Type-II EC-co-cultured SMCs were determined by immunostaining studies (Figure 4G).

Collectively, LTBP1 is a crucial target of miR-10b in “Type-II-to-Type-I” degeneration of ECs, where loss of LTBP1 protein hampers locally concentrated existence of active TGF-β at EC-SMC interaction sites, and thus, deteriorates the suppressive activities of Type-II ECs against the proliferation of SMCs.

### Trans effects of exosomes of Type-I and Type-II ECs

A wide variety of cells secret exosomes and exert diverse biological effects by conveying miRNAs(27) and proteins(28) to adjacent and distant cells. Therefore, we examined the trans effects of exosomes that were secreted by Type-I and Type-II ECs, respectively.

Firstly, we examined whether miR-10b, which is induced specifically in Type-I ECs, could be transferred to Type-II ECs *via* an exosomal route to reduce the expression levels of LTBP1 protein in Type-II ECs, and thus, convert the anti-stenotic Type-II ECs into pro-stenotic Type-I ECs. We found that treatments of Type-II ECs with “Type-I EC exosomes” reduced the expression levels of LTPB1 protein in Type-II ECs (Figure 5A) and also abolished the ability of Type-II ECs to suppress the proliferation of SMCs (Figure 5B). Hence, “Type-I EC exosomes” exert a trans effect on Type-II ECs by hampering their anti-proliferative abilities *via* miR-10b transfer to reduce the expression of LTBP1 protein, and thus, inducing “Type-II-to Type-I” conversion.

**Figure 5.**
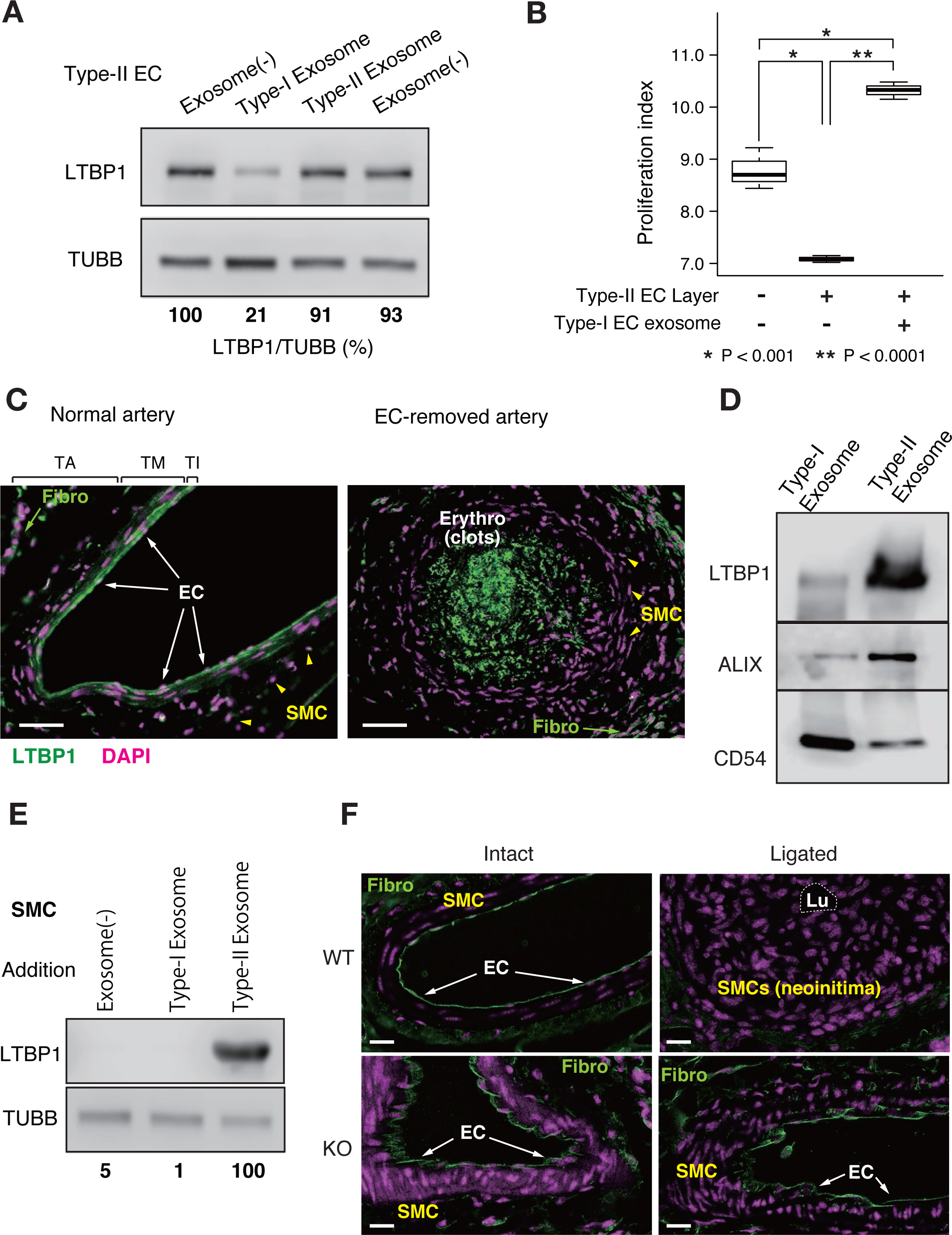
Trans-effects of EC exosomes. **A.** Western blotting for LTBP1 and TUBB using Type-II ECs treated with Type-I EC exosomes, Type-II EC exosomes or without exosomal treatments as indicated. **B.** Proliferation indexes of SMCs that were cultured on Type-II EC layers with or without additions of Type-I EC exosomes. N=3. **C.** Immunostaining for LTBP1 protein using normal femoral arteries (upper) and EC-removed femoral arteries by wire injury (WI) operations (lower). In normal arteries, LTBP1 protein (green) was continuously detected in ECs, which are lining at the luminal surface of the arteries (left, arrow heads), whereas it was sporadically detected in SMCs in tunica media (left, arrows). The strong fluorescent signals (green) detected in the lumen of the EC-removed artery are auto-fluorescence of erythrocytes in blood clots embolizing at the site of injury. Green, LTBP1; Pink, DAPI. “Fibro” indicates fibroblast in tunica adventitia. Scale=50 μm. **D**. Western blotting for LTBP1, ALIX and CD54 using the exosomes that were collected from the equivalent numbers of Type-I and Type-II ECs. **E**. Western blotting for LTBP1 and TUBB using SMCs that were treated with Type-I EC exosomes, Type-II EC exosomes or without treatments as indicated. **F.** Immunostaining for LTBP1 of the carotid arteries of wild type mice (upper) and miR-10b knockout mice (lower) that received (ligated) or did not receive (intact) the treatments for experimental atherosclerosis. Tm: tunica media; Ta: tunica adventitia. Scale=20 μm. Full-length blots are presented in Supplementary information.

Next, we examined the effects of “Type-II EC exosomes”. Since Type-II EC exosomes were negative for miR-10b expression, it does not seem possible for them to exert trans effects *via* exosomal transfer of miR-10b to adjacent cells. By contrast, Type-II EC exosomes were positive for LTBP1 protein expression, and thus, they might possibly exert trans effects *via* exosomal transfer of LTBP1 protein to strengthen TGF-β-dependent growth-inhibitory signals. To validate this hypothesis, we examined the distribution pattern of LTBP1 protein in murine arteries. ECs lining at the luminal surface of the normal arteries exclusively expressed LTPB1 protein (Figure 5C upper, arrow heads), which is consistent with the *in vitro* finding that healthy Type-II ECs expressed high levels of LTBP1 protein (Figure 4B). Although cultured SMCs were negative for LTBP1 protein expression (Supplementary Figure S2B), LTBP1 protein was sporadically detected in SMCs in tunica media (Figure 5C upper, arrows). Interestingly, LTBP1 protein became undetectable throughout the vascular walls when ECs had been mechanically removed by WI operation (Figure 5C, lower), indicating that LTBP1 protein in the vascular walls was primarily produced by ECs including the one detected in SMCs at tunica media. Since the exosomes of healthy Type-II ECs, but not those of degenerative Type-I ECs, expressed high levels of LTBP1 protein (Figure 5D), we hypothesized that LTBP1 protein was transferred from ECs to SMCs *via* an exosomal route in healthy arteries. To validate this hypothesis, we cultured SMCs with or without Type-II EC exosomes and examined whether SMCs became positive for LTBP1 protein expression. As shown in Figure 5E, SMCs that were cultured in the presence of Type-II EC exosomes, but not in the presence of Type I EC exosomes or without adding exogenous exosomes, became positive for LTBP1 protein expression. Therefore, healthy Type-II ECs, but not degenerative Type-I ECs, can transfer LTPB1 proteins to SMCs via exosomes.

Finally, we evaluated the pathological relevance of altered LTBP1 protein expressions in the development of atherosclerosis by performing immunostaining studies using the arteries of WT and miR-10b KO mice that were subjected to experimental atherosclerosis. In atherosclerotic arteries of WT mice (Figure 5C, left), LTBP1 protein expression was undetectable throughout the vascular walls (Figure 5F, upper right). By contrast, LTBP1 protein was clearly detected in ECs of the operated arteries in miR-10b KO mice (Figure 5F, lower right).

Collectively, LTBP1 proteins in vascular walls are produced by healthy ECs and transmitted to SMCs via exosomes, showing an inversely correlated expression profile with miR-10b expressions and susceptibility to atherosclerosis.

## DISCUSSION

In the current study, we unraveled the molecular events involved in the Type-I/Type-II quality control of ECs (Figure 6). In healthy arteries, ECs express LTBP1 proteins at high levels, some of which are simply secreted to extracellular spaces and some of which are transferred to SMCs *via* exosomes. Owing to abundant LTBP1 proteins that are continuously supplied by *healthy* ECs, active TGF-β exists stably in a locally concentrated manner at the site of EC-SMC interactions in tunica intima. Then, locally concentrated active TGF-β stably transmit potent growth-inhibitory signals against SMCs, thus preventing the development of neointima for the homeostatic maintenance of vascular structures. When ECs are damaged by noxious stimuli, however, Type-II ECs are degenerated into Type-I ECs. In the degenerative Type-I ECs, the expression of miR-10b is induced at the downstream of RGS5 and, as a result, the expression levels of LTBP1 protein are down-regulated. Moreover, Type-I ECs secret miR-10b^high^ exosomes to neighboring miR-10b^low^/LTBP1^high^ Type-II ECs to convert them into miR-10b^high^/LTBP1^low^ Type-I ECs. Therefore, the process of “Type-II-to-Type-I” degeneration of ECs will be accelerated and the damaged arteries will undergo stenotic changes *via* neointima formation

**Figure 6.**
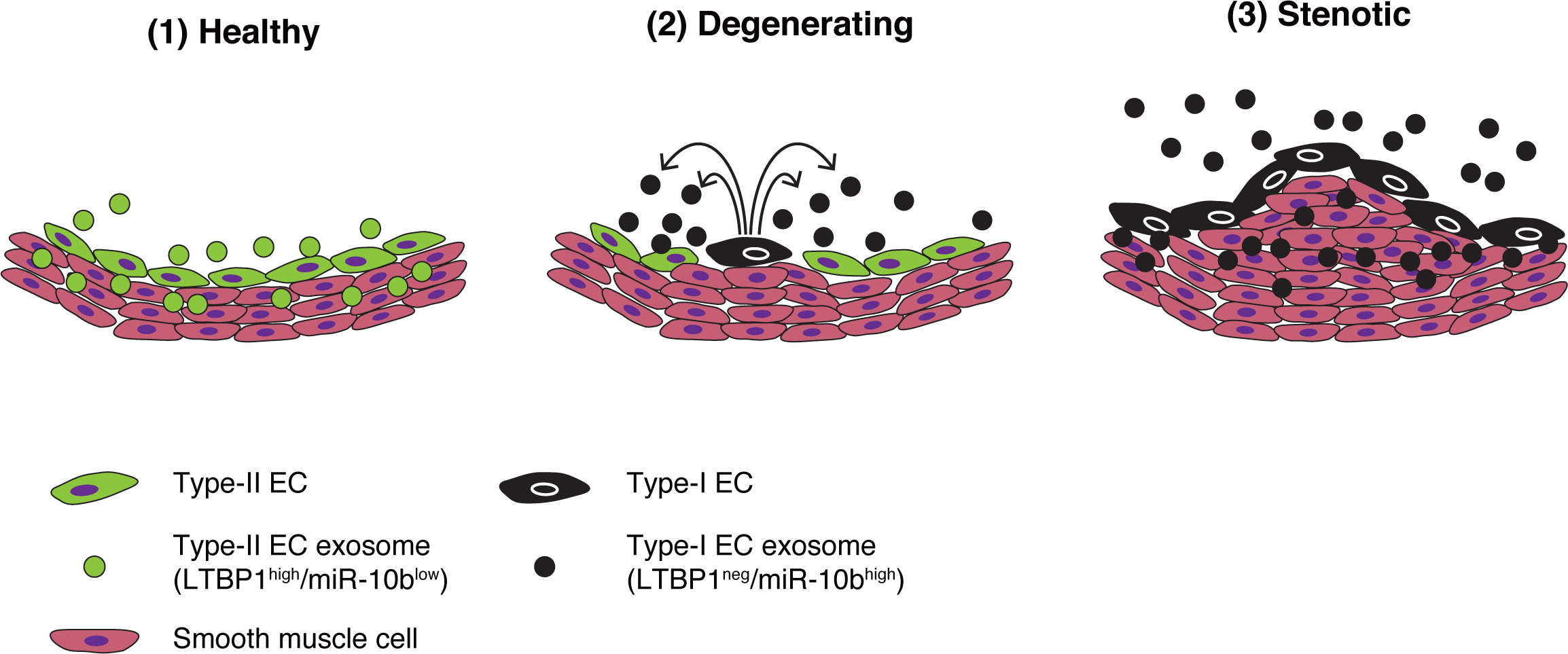
Model for the development of stenosis. (1) In healthy states, ECs retain Type-II phenotype and express LTBP1 protein at high levels. LTBP1 proteins are secreted to the extracellular matrix in tunica intima and further distributed toward tunica media. Healthy ECs also secret miR-10b^low^/LTBP1^high^ exosomes, which convey LTBP1 protein to SMCs in tunica media. By virtue of abundant LTBP1 protein, SMCs continuously receive potent TGF-β signals and the proliferations of SMCs are strongly inhibited. (2) When noxious stimuli are added to ECs, ECs are degenerated into Type-I, where miR-10b expressions are induced and LTBP1 protein expressions are reduced. Moreover, degenerative ECs secret miR-10b^high^ exosomes, which exert trans effects on adjacent Type-II ECs to convert them into Type-I ECs. Hence, the degenerating process is accelerated. (3) When noxious stimuli are continuously added to ECs, the majority of ECs at the damaged portion are inevitably converted into Type-I. As a result, LTBP1 proteins disappear from the vascular walls. Under these circumstances, TGF-β-dependent growth-inhibitory signals are waned and SMCs undergo hyper-proliferation to create neointima, thus developing stenosis of the arteries.

The molecular mechanism of Type-II EC-mediated growth inhibition of SMCs is highly analogous to the case of nonmyelinated Schwann cell (NSC)-mediated quiescence induction of HSCs in the bone marrow(12). Although TGF-β signaling is involved in the formation of stem cell niches in other cases such as melanocyte stem cells(29) and adipocyte stem cells(30), the expression level of TGF-β *per se* is the key determinant for growth inhibition of stem cells in those cases. By contrast, TGF-β protein is produced by diverse cells in the bone marrow, and thus, the expression level of TGF-β cannot be a key element for the determination of growth-suppressive potentials. Since the half-life of active TGF-β1 in the blood plasma is considerably short (ca. 2 min)(31), a certain system to stabilize TGF-β signaling is required for the formation of stem cell niches in bloodstream-rich organs such as the bone marrow. It is known that latent TGF-β1 makes a complex with LTBP1 under physiological conditions, and thus, obtain a greatly extended blood plasma half-life (> 100 min)(32). Hence, LTBP1 protein works as a crucial determinant for the formation of the NSC-based HSC niche in the bone marrow. Since ECs are constantly exposed to bloodstream, it would be highly reasonable that ECs employ LTBP1 as in the case of NSC-based HSC niche for stable implementation of their growth-inhibitory functions to protect injured arteries from undergoing ischemic changes.

Ischemic heart disease is the leading cause of death in the world. Although anticholesteremic agents have provided beneficial outcomes, there are still a number of refractory cases. For example, disseminated atherosclerotic changes are commonly detected in the coronary arteries in patients with diabetic angiopathy, which are resistant to anticholesteremic drug treatments. To develop effectual therapeutic agents for such cases, it is important to elucidate the detailed molecular basis for Type-I/Type-II quality control of ECs. The current study is grounded on our unique finding that we obtained from the analyses regarding human ES/iPS cell-derived ECs, by which the existence of anti-proliferative Type-II ECs were for the first time shown(1). Extending our previous achievement to identify RGS5 as a master gene responsible for “Type-II-to-Type-I” conversion of ECs, we have determined miR-10b as the crucial downstream mediator for quality degradation of ECs. It is reported that the miR-10b expressions in aortic blood plasma are elevated in patients of coronary artery disease with low collateral capacity(33). Our finding that miR-10b expressions in blood plasma exosomes were elevated in mice with arterial injuries (Figure 2F) is consistent with that finding. We further showed the pathological significance of miR-10b: lack of miR-10b expression effectively attenuates the development of atherosclerosis even in the presence of hypercholesterolemia. Therefore, screening miR-10b-targetting agents would be of use for the therapeutic development of ischemic diseases of various causes.

Pathological impact of miR-10b has been studied intensively in the field of cancer: miR-10b is shown to be involved in malignant progression of glioblastoma multiforme(13) and invasive breast cancer(14). It might be possible that accumulated miR-10b in the circulating blood as a result of miR-10b^high^ exosome secretion by injured ECs promotes malignant progression of cancer. Metabolic syndrome increases risk of cancer *via* multiple pathways including insulin/insulin-like growth factor (IGF) system, estrogen and pro-inflammatory cytokines(34). Our finding proposes an additional mechanism for the enhanced cancer progression in metabolic syndrome: vascular damages stimulate the emergence miR-10b^high^ pro-proliferative Type-I ECs, which then disseminate miR-10b^high^ exosomes throughout the body *via* circulating blood and thus enhance malignant progression of cancers. In this sense, miR-10b would provide a promising target for drug discovery in preventing metabolic syndrome-associated cancer progressions.

## ACKNOWLEDGMENTS

This work was supported by a Grant-in-Aid from Japan Science and Technology Agency, PRESTO (JPMJPR12M4) and the National Center for Global Health and Medicine (26-105, 26-113 and 29-1001). Authors greatly thank Mr. Shinnosuke Suzuki, Mr. Yoshinori Yanagi and Ms Rieko Yanobu-Takanashi for technical assistances.

## AUTHOR CONTRIBUTIONS

M.N. designed the study, was involved in all aspects of the experiments and wrote the manuscript; N.K. was involved in gene/protein expression analyses, statistical analyses, and wrote the manuscript; M.O was involved in gene/protein expression analyses; K.N. and T.O. generated miR-10b knockout mice; A.Y. was contributed to administrative supports; K.S designed the research, was involved in all aspects of the experiments and wrote the manuscript. All authors discussed the results and commented on the manuscript.

## METHODS

### Cells and reagents

Human umbilical vein endothelial cells (HUVEC), human aortic endothelial cells (HAEC), human microvascular endothelial cells (HMVEC) and human coronary arterial endothelial cells (HCAEC) were purchased from Dainippon Sumitomo Pharma Co., Ltd. (Osaka Japan). Human brain microvascular endothelial cells (HBVEC) and human retinal microvascular endothelial cells (HRMVEC) were purchased from Cell Systems Corp. (Kirkland, WA, USA). All VECs were cultured using EGM™-2 BulletKit™ or EGM™-2MV BulletKit™ (Lonza Group Ltd., Basel, Switzerland). Human aortic smooth muscle cells (SMC) were purchased from Lonza Group Ltd and cultured using SmGM™-2 BulletKit™ (Lonza Group Ltd.). Cells were re-seeded at split ratios of 1:3 ~ 1:4 twice a week. Human ESC lines (KhES-1, -3) were established by the Institute for Frontier Medical Science, Kyoto University(35). The human SeV-hiPSCs(36) were established from HUVEC or BJ fibroblast. Differentiation of human ES/iPS cells into ECs was performed as previously described(37–39). Azidothymidine (A2169, Sigma-Aldrich Co. LLC, St. Louis, MO, USA) was dissolved by H_2_O. A TGFB inhibitor (LY2157299, WAKO Pure Chemical Industries, Osaka, Japan) were dissolved by DMSO. As an inhibitor for miR-10b, MISSION^®^ Lenti microRNA Inhibitor, Human, hsa-miR-10b-5p (HLTUD0044, Sigma-Aldrich Co. LLC) along with control vectors (HLTUD001C, and HLTUD002C, Sigma-Aldrich Co. LLC) was used.

### Evaluations of SMC proliferation

Proliferation of EC-co-cultured SMCs was quantitatively evaluated as previously reported(1). Briefly, ECs were γ-irradiated (40 Gy) and stained with carboxyfluorescein diacetate, succinimidyl ester (Cayman Chemical Co., Ann Arbor, MI) and SMCs were stained with PKH26 (Sigma-Aldrich Co. LLC). ECs were seeded at the density of 2×10^5^ cells/well on 0.1% gelatin-coated 24-well plates, and on the following day, SMCs were seeded at the density of 3.75×10^3^ cells/well on EC layers or gelatin layers. After 4 days, cells were harvested and subjected to flow cytometry analyses by FACSCaliburTM (BD Biosciences, San Jose, CA) and FL1 and FL2 fluorescence intensities were measured by CellQuest™ Pro software (BD Biosciences). Proliferation indexes of SMCs were calculated by statistically analyzing the fluorescence histograms of the gated FL2 (PKH26)-positive cells using ModFit LT™ software (Verity Software House Inc., Topsham, ME).

### miRNA Microarray

Total RNAs were isolated by using miRNeasy Mini Kit (Cat# 217004, Qiagen, Hilden, Germany) and subjected to Agilent human miRNA Microarray (8×60K) miRBase 19.0 (Agilent Technologies, Santa Clara, CA) by Chemicals Evaluation and Research Institute (Tokyo, Japan). Differences in miRNA expression profiles between the two groups (Type-I EC groups *versus* Type-II EC groups) were estimated as significant if P values were less than 0.01.

### Quantitative Reverse Transcription Polymerase Chain Reaction (qRT-PCR)

Total RNAs were isolated by using miRNeasy Mini Kit and complementary DNA was prepared from 1 μg of RNA using SuperScript™III First-Strand Synthesis System Kit (Thermo Fisher Scientific Inc. Waltham, MA USA). StepOnePlus™ PCR machine (Thermo Fisher Scientific Inc.) was used for qPCR by applying either FAST SYBR^®^ Green Master Mix (Thermo Fisher Scientific Inc.) with primers (TGFB1, Fw: 5’CGTGGAGGGGAAATTGAGGG3’ and Rv: 5’CCGGTAGTGAACCCGTTGATG3’; LTBP1, Fw: 5’TGCTGTCTGTATGGAGAGGC3’ and Rv: 5’CTGAAGTCAACCAAGGCGTC3’; GAPDH, Fw: 5’-CCACTCCTCCACCTTTGAC-3’and Rv:5’-ACCCTGTTGCTGTAGCCA-3’) or TaqMan™ Assays (ID: 002218 for hsa-miR-10b; ID: 001001 for RNU24; ID: 000389 for hsa-miR-15a). To normalize cytoplasmic miRNA expressions, RNU24 was used for an internal control as recommended by Applied Biosystems^®^ (Thermo Fisher Scientific Inc. http://bioinfo.appliedbiosystems.com/genome-database/gene-expression.html). To normalize exosomal miRNA expressions, miR-15a was used for an internal control as reported elsewhere^43^.

### Antibodies

Western blotting studies were performed as previously described(1) using an anti-human LTBP1 antibody (22065-1-AP) (Proteintech Group, Chicago IL, USA), anti-human LTBP2 antibody (17708-1-AP, Proteintech Group), anti-human LTBP3 (H-210) antibody (sc-98276, Santa Cruz Biotechnology Inc., Santa Cruz, CA), anti-human LTBP4 (H-293) antibody (sc-33144, Santa Cruz Biotechnology Inc.), anti-human TSG101 antibody (ab83, Abcam plc, Cambridge, UK), anti-human ALIX antibody (2171S, Cell Signaling Technology, Inc., Danvers, MA), anti-human CD54 antibody (10020-1-AP, Proteintech Group) and anti-human β-tubulin antibody (ab134185, Abcam plc). The signal detection procedure was performed using either SuperSignal West Dura (Thermo Fisher Scientific Inc.) or SuperSignal West Femto (Thermo Fisher Scientific Inc.) along with development on Hyperfilm ECL (GE Healthcare, Little Chalfont, UK) or image scanning using C-DiGit^®^ Blot Scanner for Chemiluminescent Western Blots (LI-COR, Inc. Lincoln, Nebraska USA). Immunostaining studies were performed as previously described^1^ using an anti-human TSG101 antibody (ab83, Abcam plc) or anti-SMAD3 (C67H9) antibody (Cell Signaling Technology Inc., Danvers, MA, USA). Photomicrographs were taken either by using Olympus BX51 fluorescence microscope (Olympus Optical Co. Ltd.) equipped with DP-2 TWAIN digital camera system (Olympus Optical Co. Ltd.) and cellSens^®^ standard imaging software (Olympus Optical Co. Ltd.) or BZ-X710 all-in-one fluorescence microscopy series (Keyence Corp, Osaka, Japan). For flow cytometry, cells were perforated by using Fix&Perm Cell Permeabilization Kit (GAS003) (Thermo Fisher Scientific Inc.), stained by an Alexa Fluor^®^ 647 Anti-Smad2 (pS465/pS467)/Smad3 (pS423/pS425) (BD562696, BD Bioscience) and analyzed by using FACSCalibur™ (BD Bioscience, Franklin Lakes, NJ, USA) as previously described(1).

### Transfection

Transfection using simian immunodeficiency virus (SIV)-based RGS5 expression vector (ID Pharma Co., Ltd., Ibaraki, Japan) was performed as previously described(1). An expression vector for hsa-miR-10b (Cat# SC400025, OriGene Technologies Inc., Rockville, MD) or a control pCMVMIR vector (OriGene Technologies Inc.) was introduced by using Nucleofector™ (Lonza Group Ltd., Basel, Switzerland) with Amaxa HUVEC Nucleofector Kit (#VPB-1002, Lonza Group Ltd.) for primary cultured VECs and Amaxa Basic Nucleofector Kit Primary Endothelial Cells (#VPI-1001, Lonza Group Ltd.) for hESdECs and hiPSdECs. A lentiviral vector-based miR10b inhibitor (Cat# HLTUD0044 in MISSION^®^ Lenti microRNA Inhibitor, Sigma-Aldrich Co. LLC.) was transfected according to the manufacturer’s guidance (MOI=50).

### Exosome preparation

Exosomes were prepared from either cell culture supernatant using miRCURY Exosome Isolation Kit Cells, Urine and CSF (Cat# 300102, Exiqon A/S, Vedbaek, Denmark) or murine sera using miRCURY Exosome Isolation Kit Serum and plasma (Cat# 300101, Exiqon A/S) according to the manufacture’s guidance.

### Animal experiments

Wire injury (WI) operation was performed in the femoral artery as previously described(4). Experimental atherosclerosis was performed as previously reported(36). Briefly, five week-old mice were fed with Paigen Diet containing 1.25% Cholesterol, 0.5% Sodium Cholic Acid, 35 kcal% FAT (D12336, Research Diets, Inc., New Brunswick, NJ, USA). At six weeks of age, they were subjected to partial carotid ligation by ligating the three branches (external carotid artery, internal carotid artery and occipital artery) of the left common carotid artery (CLA). At nine weeks of age, the conditions of CLAs were ultrasonically examined by using Vevo^®^2100 (Primetech Corp., Tokyo, Japan). Mice were then euthanatized and CLAs were removed for histological examinations.

CRISPR/Cas9-mediated genome editing in mice was performed as described previously(40) with some modifications. Briefly, single-guide (sg) RNA expression vector for target sequence with a T7 promoter was synthesized, and transcribed *in vitro* using MEGAshortscript Kit (Life Technologies, Carlsbad, CA). *hCas9* mRNA was synthesized using mMESSAGE mMACHINE T7 kit (Life Technologies), and was polyadenylated with polyA tailing kit (Life Technologies). Introduction of purified sgRNA, *hCas9* mRNAs into fertilized eggs from C57BL/6NCr (Japan SLC Inc., Hamamatsu, Japan) by electroporation was performed as described previously^(41)^. After the electroporated oocytes were cultured overnight *in vitro,* two-cell embryos were transferred into the oviducts of pseudopregnant ICR females (CLEA Japan Inc., Tokyo, Japan). Tail genomic DNA of offsprings was isolated using standard methods. The genomic sequence at miR-10b gene locus was amplified by PCR using a forward primer (5’-ACCATCTCTGAAAGCCAGGC-3’) and a reverse primer (5’-TGGAGCTTTCTGGCCTTTCA-3’). The mutation was confirmed by Sanger sequencing.

All mice were housed in air-conditioned animal rooms at an ambient air temperature of 22 ± 2ºC and relative humidity of 50 ± 15%, under specific pathogen-free conditions with a 12-h light/dark cycle (8:00 to 20:00 light period). They were fed a standard diet (CE-2, CLEA Japan, Inc., Tokyo Japan) with free access to drinking water. All animal experiments were approved by the Animal Care and Use Committee of the National Center for Global Health and Medicine (NCGM) Research Institute (Permission Numbers: 16014, 16015, 16040, 17017 and 17021) and conducted in accordance with institutional procedures.

### Statistical analyses

All statistical analysis and drawing graphs were performed by each function in R software (ver 3.0.1)(42). One-sample t-test was performed for Fig. 1E and 2A, and Welch’s Two-sample t-test was used for Fig. 1C, D, Fig. 2B, C, D and Fig. 3A. To do multiple comparison (for Fig. 2E, Fig. 3C, Fig. 4E, F and Fig. 5B), Tukey’s HSD (honest significant difference) test was used, following to Bartlett test for checking of their homogeneity of variances. Box plot (Fig. 1C, 1D, 1E, Fig. 2, Fig. 5B) were made by the ‘boxplot’ function. Box indicates the value from the 1^st^ and 3^rd^ quartile (interquartile range: IQR) and a thick line in the box shows a median. The upper and lower whisker show the maximum and minimum values, respectively, which do not exceed 3/2 IQR beyond the box. Bar plot (Fig. 4E and 4F) were drawn by the ‘barplot’ function. Error bar indicated ±SD.

### Data availability

Raw data of miRNA Microarray can be accessed at the Gene Expression Omnibus (GEO) under accession ID: GSE102207.

